# Lithium inhibits tryptophan catabolism via the inflammation-induced kynurenine pathway in human microglia

**DOI:** 10.1101/2020.11.24.388470

**Authors:** Ria Göttert, Pawel Fidzinski, Larissa Kraus, Ulf Christoph Schneider, Martin Holtkamp, Matthias Endres, Karen Gertz, Golo Kronenberg

**Affiliations:** Charité-Universitätsmedizin Berlin, corporate member of Freie Universität Berlin, Humboldt-Universität zu Berlin, and Berlin Institute of Health, Klinik für Neurologie und Abteilung für Experimentelle Neurologie, Berlin, Germany; Center for Stroke Research Berlin (CSB), Berlin, Germany; Epilepsy-Center Berlin-Brandenburg, Berlin, Germany; NeuroCure Cluster of Excellence, Berlin, Germany; Charité-Universitätsmedizin Berlin, corporate member of Freie Universität Berlin, Humboldt-Universität zu Berlin, and Berlin Institute of Health, Klinik für Neurochirurgie, Berlin, Germany; German Center for Neurodegenerative Diseases (DZNE), partner site Berlin, Germany; German Centre for Cardiovascular Research (DZHK), partner site Berlin, Germany; College of Life Sciences, University of Leicester, United Kingdom; Leicestershire Partnership National Health Service Trust, Leicester, United Kingdom

**Keywords:** depression, kynurenine, lithium, microglia, serotonin, tryptophan

## Abstract

Activation of the kynurenine pathway may lead to depletion of the serotonin precursor tryptophan, which has been implicated in the neurobiology of depression. This study describes a mechanism whereby lithium inhibits inflammatory tryptophan breakdown. Upon activation, immortalized human microglia showed a robust increase in indoleamine-2,3-dioxygenase (*IDO1*) mRNA transcription, IDO1 protein expression, and activity. Further, chromatin immunoprecipitation verified enriched binding of both STAT1 and STAT3 to the *IDO1* promoter. Lithium counteracted these effects, increasing inhibitory GSK3β^S9^ phosphorylation and reducing STAT1^S727^ and STAT3^Y705^ phosphorylation levels in activated cells. Experiments in primary human microglia and human induced pluripotent stem cell (hiPSC)-derived microglia corroborated lithium’s effects. Moreover, IDO activity was reduced by GSK3 inhibitor SB-216763 and STAT inhibitor nifuroxazide via downregulation of P-STAT1^S727^ and P-STAT3^Y705^. Our study demonstrates that lithium inhibits the inflammatory kynurenine pathway in the microglia compartment of the human brain.

## Introduction

The monovalent cation lithium has been in medical use for decades. Lithium remains a mainstay in the treatment and prophylaxis of bipolar disorder (Geddes et al., 2004; Malhi et al., 2017). Lithium is also effective in the prophylaxis of severe unipolar major depression (Tiihonen, 2017), and adjunctive lithium constitutes a promising therapeutic approach for patients who do not respond adequately to antidepressants (e.g. (Bauer et al., 2010)). Moreover, numerous reports have found that lithium exerts anti-suicidal effects in patients with mood disorders (reviewed in (Smith and Cipriani, 2017)).

Immunological mechanisms and, in particular, microglia activation, seem to play an important role in the etiopathogenesis of affective disorders in human patients and social-defeat induced behavioral changes in rodents (e.g. (Jeon and Kim, 2016; Kopschina Feltes et al., 2017; McKim et al., 2018; Nie et al., 2018; Torres-Platas et al., 2014; Yirmiya et al., 2015)). Upregulation of indoleamine-2,3-dioxygenase (IDO1), the rate-limiting enzyme converting tryptophan into kynurenine in activated microglia, may underlie reduced availability of serotonin in neuroinflammation. Anti-inflammatory effects of lithium have been reported by a number of recent studies (e.g. (Ajmone-Cat et al., 2016; Cao et al., 2017; Dong et al., 2014)). Moreover, numerous case reports have documented an increased risk of serotonin syndrome precipitated by co-prescription of lithium with antidepressants (e.g., (Adan-Manes et al., 2006; Prakash et al., 2017; Sobanski et al., 1997)). A study in rats treated with chronic lithium found increased levels of tryptophan alongside increased concentrations of serotonin metabolite 5-hydroxyindole acetic acid (5-HIAA) in brain (Perez-Cruet et al., 1971). Notwithstanding these observations, the effect of lithium on kynurenine pathway activity has not been investigated prior to this study. Using immortalized microglia, primary adult human microglia cultured from surgical specimens, and human iPSC-derived microglia-like cells, we herein confirmed the hypothesis that lithium decreases IDO1 activity in activated microglia and studied the underlying intracellular signaling pathways.

## Results

### Lithium reduces microglial expression of IDO1 by inhibiting GSK3β, preventing STAT1/3 activation

To examine mRNA transcription of the first rate-limiting kynurenine pathway enzyme in response to TLR4 and type II interferon signaling, we treated a commercially available human microglia cell line with lipopolysaccharide (LPS) and IFN-γ. We found that IFN-γ, but not LPS, was able to activate immortalized human microglia, inducing a robust *IDO1* mRNA increase which was significantly reduced when lithium was added to the cultures (Figure 1A). Importantly, lithium (10 mM) did not exert toxic effects on microglia (Figure S1B). In contrast to *IDO1*, *IDO2* and *TDO* were not induced by IFN-γ treatment in immortalized human microglia (Figure S1A). To quantify intracellular protein levels, we performed a Western blot against IDO1 and actin beta (ACTB). IDO1 was only detectable after IFN-γ treatment, and the IDO1/ACTB ratio was significantly decreased (~ 60%) in the presence of lithium (Figure 1B). Kynurenine (KYN) release into the cell culture medium was strongly increased by IFN-γ activation. Treatment with lithium significantly reduced this effect (Figure 1C). Tryptophan (TRP) degradation increased massively after IFN-γ activation while co-treatment with lithium reduced tryptophan catabolism. The ratio of KYN/TRP is an indirect measure of IDO activity. IDO activity was significantly reduced when IFN-γ stimulated immortalized human microglia were simultaneously incubated with lithium (Figure 1C). Next, we aimed to investigate the pathway that might regulate IFN-γ induced *IDO1* transcription in human microglia cells. Overall, 19 signaling molecules involved in the regulation of immune and inflammatory responses were studied (Figure 1D). Three of the 19 signaling molecules studied showed strong induction in response to IFN-γ treatment (P-STAT1^Y701^, P-STAT1^S727^, P-STAT3^Y705^), with lithium significantly reducing STAT1^S727^ and STAT3^Y705^ phosphorylation levels in IFN-γ treated cells (Figure 1D, E). Western blots against phosphorylated STAT1^S727^ and STAT3^Y705^ further corroborated this finding (Figure 1F). Moreover, lithium increased GSK3β^S9^ phosphorylation (Figure 1G), resulting in inhibition of GSK3β activity (Jope and Johnson, 2004). Next, we performed chromatin immunoprecipitation to assess binding of STAT1 and STAT3 to the *IDO1* promoter. Primers spanning the STAT1, STAT3 binding site (−1086 bp relative to transcription start site according to Eukaryotic Promotor Database EPD, JASPAR CORE vertebrates) were used. Enriched binding of both STAT1 and STAT3 to the *IDO1* promoter was observed after IFN-γ activation of immortalized human microglia cells (Figure 1H).

**Figure 1.**
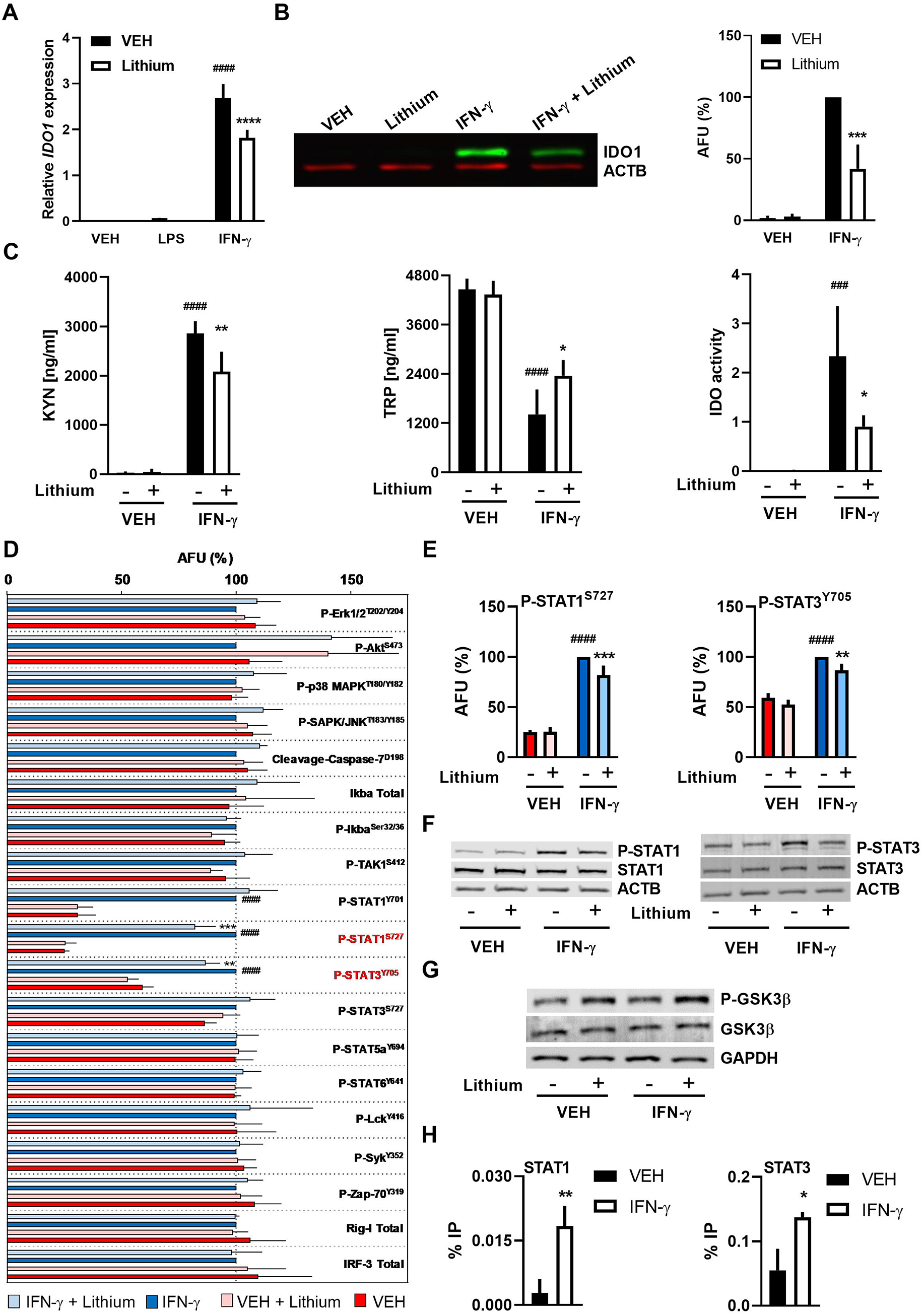
Lithium suppresses IFN-γ induction of IDO1 in immortalized human microglia by inhibiting STAT1 and STAT3 signaling. **(A)***IDO1* mRNA expression was measured after 12 h treatment with IFN-γ. N=4 independent experiments, ^####^p<0.0001 vehicle vs. IFN-γ, ****p<0.0001 IFN-γ vs. IFN-γ and lithium, two-way ANOVA followed by Tukey’s post hoc test. (**B)** Representative Western Blot analysis and relative fluorometric quantification after 12 h treatment with IFN-γ. AFU, arbitrary fluorescent units. Values are expressed as percentage of the IFN-γ-treatment condition (100%). N=3 independent experiments, ***p<0.001 IFN-γ vs. IFN-γ and lithium, two-way ANOVA followed by Tukey’s post hoc test. **(C)** The concentrations of kynurenine (KYN) and tryptophan (TRP) were measured after 24 h treatment with IFN-γ. The KYN/TRP ratio was used as a measure of IDO activity. N=4 independent experiments, ^###^p<0.001, ^####^p<0.0001 vehicle vs. IFN-γ, *p<0.05, **p<0.01 IFN-γ vs. IFN-γ and lithium, two-way ANOVA followed by Tukey’s post hoc test. **(D, E)** After 1 h treatment with IFN-γ and/or lithium, an intracellular cell signaling antibody array (PathScan®) comprising a panel of 19 signaling molecules was performed. Lithium significantly reduced IFN-γ-induced phosphorylation of STAT1 at serine 727 as well as phosphorylation of STAT3 at tyrosine 705. N=5 independent experiments, ^####^p<0.0001 for the comparison of IFN-γ vs. vehicle, **p<0.01, ***p<0.001 for the comparison IFN-γ vs. IFN-γ and lithium. Two-way ANOVA followed by Tukey’s post hoc test. **(F)** Representative Western blots confirming the results of the PathScan® analysis. **(G)** Lithium increases phosphorylation of GSK3β. Cells were harvested after 1 h treatment with IFN-γ and/or lithium. **(H)** Real-time PCR analysis of *IDO1* promoter fragments recovered in chromatin immunoprecipitation (ChIP) experiments with antibodies against STAT1 and STAT3. Chromatin was prepared after 3 h treatment with IFN-γ. Quantitative PCR data are presented as percent of input (% IP). N=3, unpaired t-test, *p<0.05, **p<0.01.

### Lithium reduces kynurenine pathway activity in activated primary human microglia

Since immortalized human microglia proved insensitive to LPS, *IDO1* transcription was further investigated in primary human microglia. These primary cells were cultured from cortical brain biopsies of patients suffering from temporal lobe epilepsy. Cultures expressed typical microglia markers such as IBA1, PU.1, TMEM119, CD45, and P2YR12 (Figure S2A). Moreover, these cells phagocytosed bacterial particles and retained a rod-shaped morphology even after activation with IFN-γ or LPS (Figure S2B, C)), which is in contrast to the amoboid shape observed after LPS treatment in postnatal primary mouse microglia (Bronstein et al., 2013; Tam and Ma, 2014). LPS and IFN-γ (Figure 2A) were both found to be strong inducers of *IDO1* mRNA transcription in primary human microglia. Lithium co-treatment significantly reduced this effect (Figure 2A). MTT assay confirmed that lithium was non-toxic to primary human microglia at the concentration of 10 mM (Figure S1D). There was no apparent regulation of either *IDO2* or *TDO* in primary human microglia in response to stimulation with LPS or IFN-γ, the levels of both transcripts being just above the detection limit (Figure S1C). To evaluate IDO1 protein expression in primary human microglia, we performed immunocytochemistry and measured fluorescence intensities. IDO1 became detectable after stimulation of primary human microglia with either LPS or IFN-γ. Again, co-treatment with lithium reduced IDO1 expression (Figure 2B). Furthermore, IDO activity in primary human microglia was investigated following treatment with LPS or IFN-γ (Figure 2C). The kynurenine/tryptophan ratio was studied in primary cells derived from 4 different individuals. While LPS and IFN-γ increased IDO activity, the addition of lithium to the cultures significantly reduced this effect (Figure 2C). Since lithium has well-known anti-inflammatory properties (Huang et al., 2009; Nahman et al., 2012; Yuskaitis and Jope, 2009), we also measured the expression of anti-inflammatory cytokine IL-10. We found IL-10 to be specifically induced in microglia after LPS treatment. Co-treatment with lithium caused a massive increase in IL-10 levels of more than four times the levels observed in LPS-stimulated cells (Figure 2D). In a next step, we investigated the phosphorylation status of transcriptional regulators STAT1 und STAT3 (Figure 2E). Given the limited availability of primary cells, the expression of STAT1 and STAT3 was close to the Western blot detection limit in our experiments. However, primary human microglia showed a clear increase in phosphorylation of STAT1^S727^ after activation with LPS (Figure 2E, top). IFN-γ activation increased phosphorylation of both STAT1^S727^ and STAT3^Y705^ (Figure 2E, top and center). Furthermore, dephosphorylation of STAT1^S727^ and STAT3^Y705^ was clearly apparent after lithium treatment of IFN-γ activated primary human microglia (Figure 2E). Finally, GSK3β is constitutively expressed at a high level in primary human microglia. Again, lithium increased inhibitory phosphorylation at serine 9 (Figure 2E, bottom).

**Figure 2.**
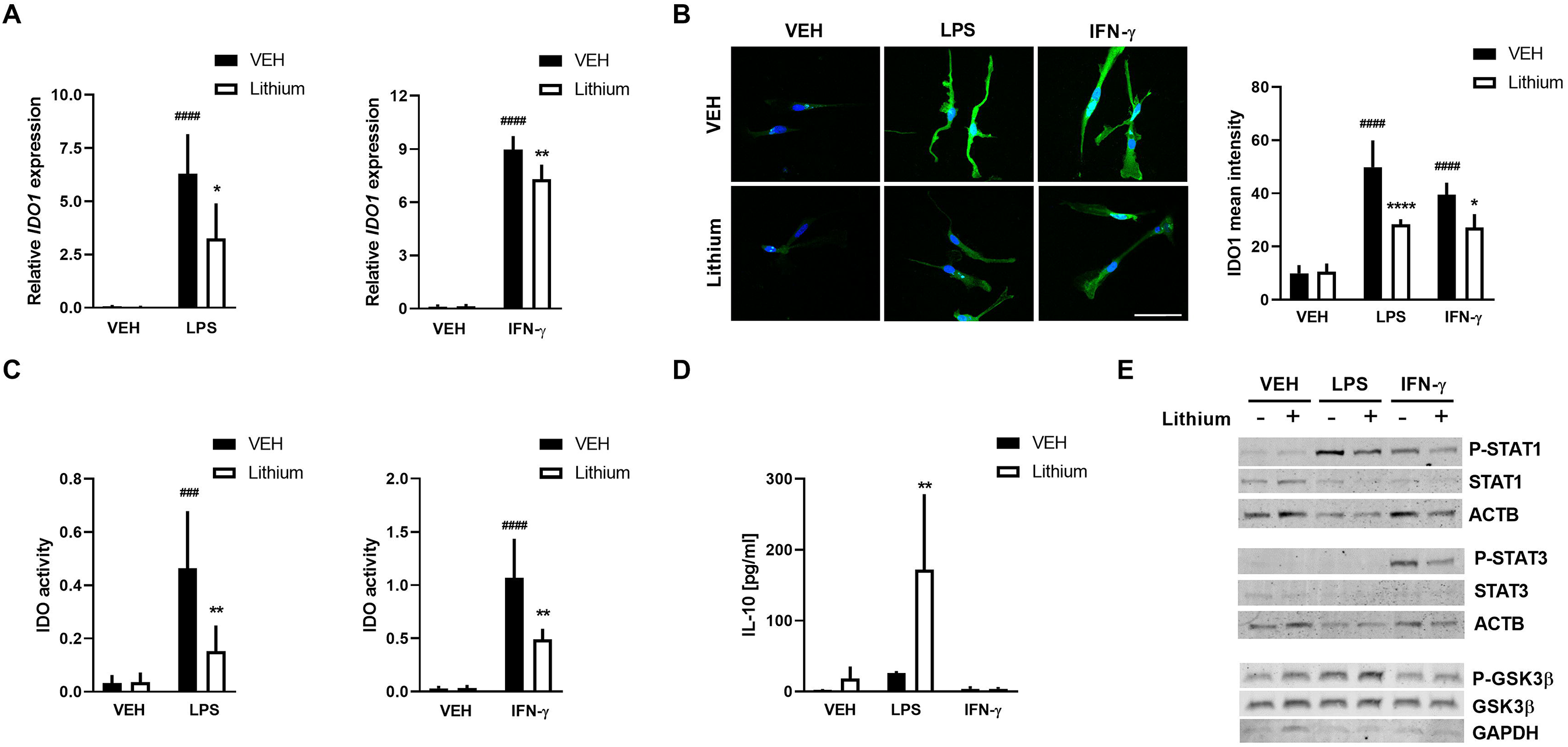
Lithium reduces kynurenine pathway activity in activated primary human microglia. **(A)** *IDO1* mRNA expression was measured after 24 h stimulation with LPS or IFN-γ. N=4 independent experiments, ^####^p<0.0001 vehicle vs. stimulation condition, *p<0.05, **p<0.01 stimulation condition vs. stimulation condition with lithium, two-way ANOVA followed by Tukey’s post hoc test. **(B)** Representative immunofluorescent images demonstrating that IDO1 protein expression (green) is increased 24 h after treatment with either LPS (upper middle image) or IFN-γ (upper right image). The bottom images show that lithium reduced this increase in IDO1 in activated microglia. Blue, Hoechst nuclear counterstain. Green fluorescence intensity (i.e., IDO1 expression) was quantified using Zen® software. Microglia from a single surgical specimen were used. Five images containing between 2 to 11 microglia cells each were analyzed per condition. Mean fluorescence intensity was calculated for each image (n=5), ^####^p<0.0001 vehicle vs. stimulation condition, *p<0.05 IFNγ vs. IFNγ and lithium, ****p<0.0001 LPS vs. LPS and lithium, two-way ANOVA followed by Tukey’s post hoc test. **(C)** IDO1 activity was quantified as KYN/TRP ratio 24 h post-treatment. N=5 independent experiments, ^###^p<0.001, ^####^p<0.0001 vehicle vs. stimulation condition, **p<0.01 for the effect of lithium in stimulated microglia. Two-way ANOVA followed by Tukey’s post hoc test. **(D)** IL-10 protein concentration was measured in cell culture supernatant 24 h after treatment. N=4 independent experiments, **p<0.01 LPS vs. LPS and lithium, two-way ANOVA followed by Tukey’s post hoc test. **(E)** Representative Western blots confirming the results obtained in immortalized human microglia. Note that, unlike immortalized human microglia, primary human microglia can be activated by both LPS and IFN-γ. Cells were harvested 1 h post-treatment.

### Inhibitors of GSK3β and STAT are able to modulate IDO activity in immortalized and primary human microglia

Next, we investigated whether modulation of GSK3β and STAT activity alters microglia function and IDO activity. SB-216763 (SB-21) was used to inhibit GSK3β activity (Coghlan et al., 2000). Nifuroxazide (Nz) was used to inhibit STAT signaling (Nelson et al., 2008). MTT assays confirmed that, at the concentrations used in subsequent experiments, inhibitors did not affect cell viability of immortalized (Figure 3A) or primary human microglia (Figure 3B). Activity measurements revealed that both SB-216763 and nifuroxazide significantly reduced IDO activity in immortalized human microglia stimulated with IFN-γ (Figure 3C). IDO1 protein expression in IFN-γ activated immortalized cells was significantly reduced after inhibitor treatment (Figure 3D). To further elucidate the effects of SB-216763 and nifuroxazide on the regulation of IDO1, we investigated whether phosphorylation of P-STAT1^S727^ and STAT3^Y705^ would be modulated by either compound following stimulation with LPS or IFN-γ. The results of Western blots indicated that treatment with SB-216763 reduced P-STAT1^S727^ and P-STAT3^Y705^. Treatment with nifuroxazide inhibited P-STAT3^Y705^ (Figure 3D). Finally, in primary human microglia, inhibitor treatment of LPS or IFN-γ activated cells resulted in significantly reduced IDO1 protein expression (Figure 3E) and reduced IDO activity (Figure 3F).

**Figure 3.**
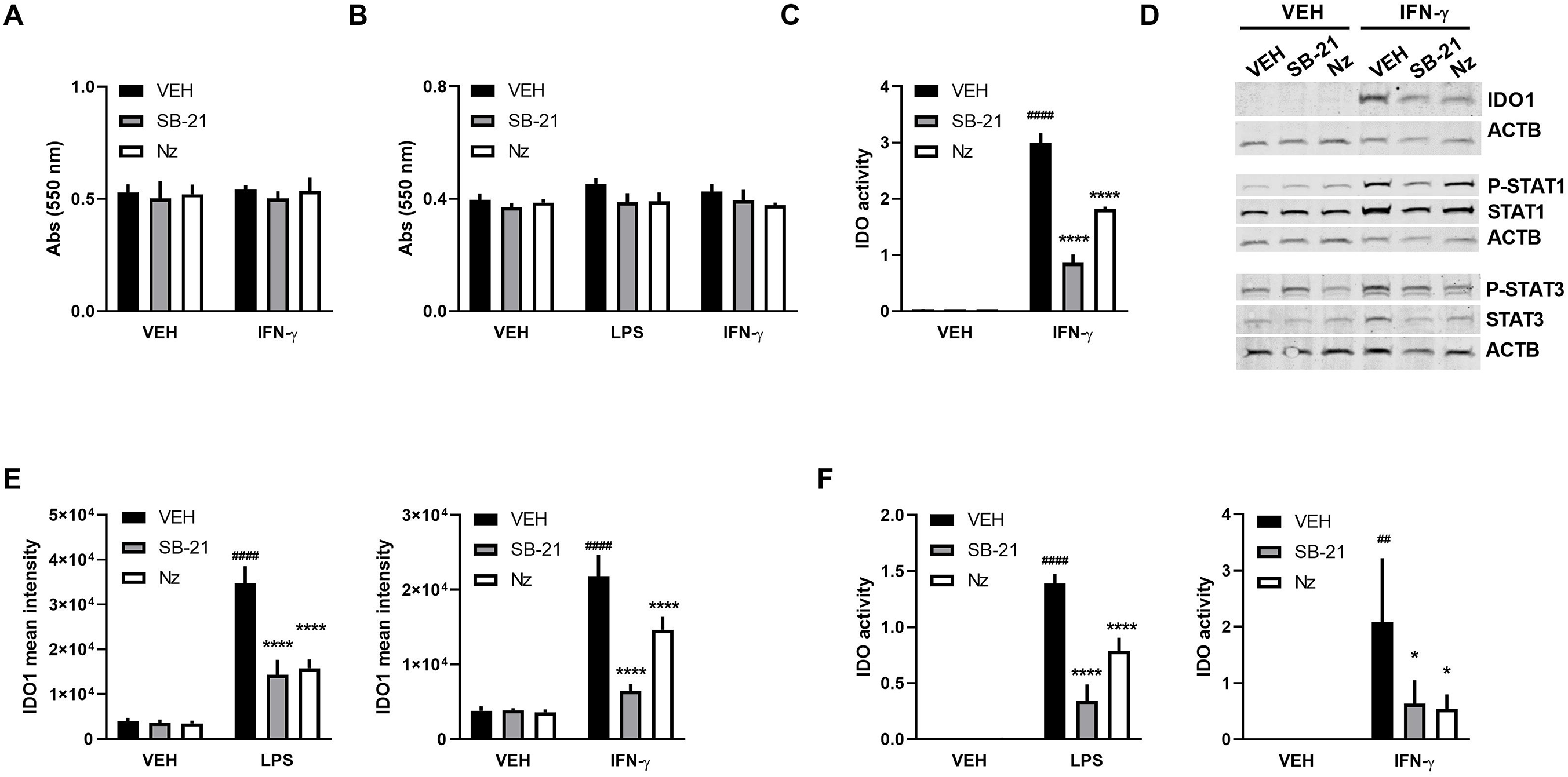
Inhibitors of GSK3β and STAT are able to modulate IDO activity in immortalized and primary human microglia. **(A, B)** Cell viability following 24 h treatment with GSK3 inhibitor SB-216763 (SB-21), STAT inhibitor nifuroxazide (Nz), or vehicle was studied using MTT assay. **(A)** Immortalized human microglia **(B)** primary human microglia. N=3. **(C)** HPLC analysis of cell culture supernatant to assess IDO activity in immortalized human microglia. Analyses were performed 24 h after stimulation with IFN-γ and co-treatment with either SB-21, Nz, or vehicle. N=3, two-way ANOVA followed by Tukey’s post hoc test. ^####^p<0.0001 vehicle vs. IFN-γ, ****p<0.0001 IFN-γ vs. IFN-γ and co-treatment with either inhibitor. **(D)** Western blot of cell extracts of immortalized human microglia. IDO1 expression was studied 12 h after IFN-γ treatment. STAT1, STAT3 as well as phosphorylation of STAT1 at serine 727 and phosphorylation of STAT3 at tyrosine 705 were assessed following 1 h IFN-γ stimulation. Actin beta (ACTB) was used as a loading control. **(E)** IDO1 expression in primary human microglia was quantified based on fluorescence intensity. N=5, two-way ANOVA followed by Tukey’s post hoc test ^####^p<0.0001 vehicle vs. LPS / IFN-γ, ****p<0.0001 LPS / IFN-γ vs. LPS / IFN-γ and co-treatment with an inhibitor **(F)** HPLC analysis of cell culture supernatant to assess IDO activity in primary human microglia. N=3, two-way ANOVA followed by Tukey’s post hoc test. ^####^P<0.0001 vehicle vs. LPS, ^##^p<0.01 vehicle vs. IFN-γ, ****p<0.0001 LPS vs. LPS and co-treatment with either inhibitor *p<0.05 IFN-γ vs. IFN-γ and co-treatment with either inhibitor.

### Tryptophan metabolism in hiPSC-derived microglia

Considering the limited availability of human primary cells, we also studied tryptophan metabolism in hiPSC-derived microglia using three different human iPSC lines. Human iPSC-derived microglia were differentiated according to an existing protocol (Abud et al., 2017). This method produces pure cultures with characteristic rod-shaped cells (Figure 4A). These cells phagocytose bacterial particles (Figure 4B) and express typical microglia markers such as CXCR1, IBA1, P2YR12, PU.1, and CD45, but not TMEM119 (Figure 4C). IDO1 immunostaining showed that all cells expressing CD45 after IFN-γ stimulation also expressed the IDO1 protein. By contrast, IDO1 immunoreactivity was not found in all cells after LPS activation (Figure 4D). LPS treatment of hiPSC-derived microglia resulted in activation of cytokines IL-10, TNF-α, IL-6, IL-8, IL-1β, and IL-12p70 (Figure 4E). The significant upregulation of IL-10 in LPS activated primary cells in the presence of lithium was recapitulated in hiPSC-derived microglia (Figure 2D and Figure 4E). Gene expression analyses of tryptophan degrading enzymes *IDO1*, *IDO2*, and *TDO* were performed. Stimulation with LPS or IFN-γ resulted in strong upregulation of *IDO1* mRNA transcription with no significant effects on *IDO2* or *TDO* (Figure 4F). Finally, lithium treatment decreased *IDO1* mRNA transcription after LPS or IFN-γ activation (Figure 4G). Increased IDO activity was found after IFN-γ activation, an effect which was again attenuated by lithium (Figure 4H).

**Figure 4.**
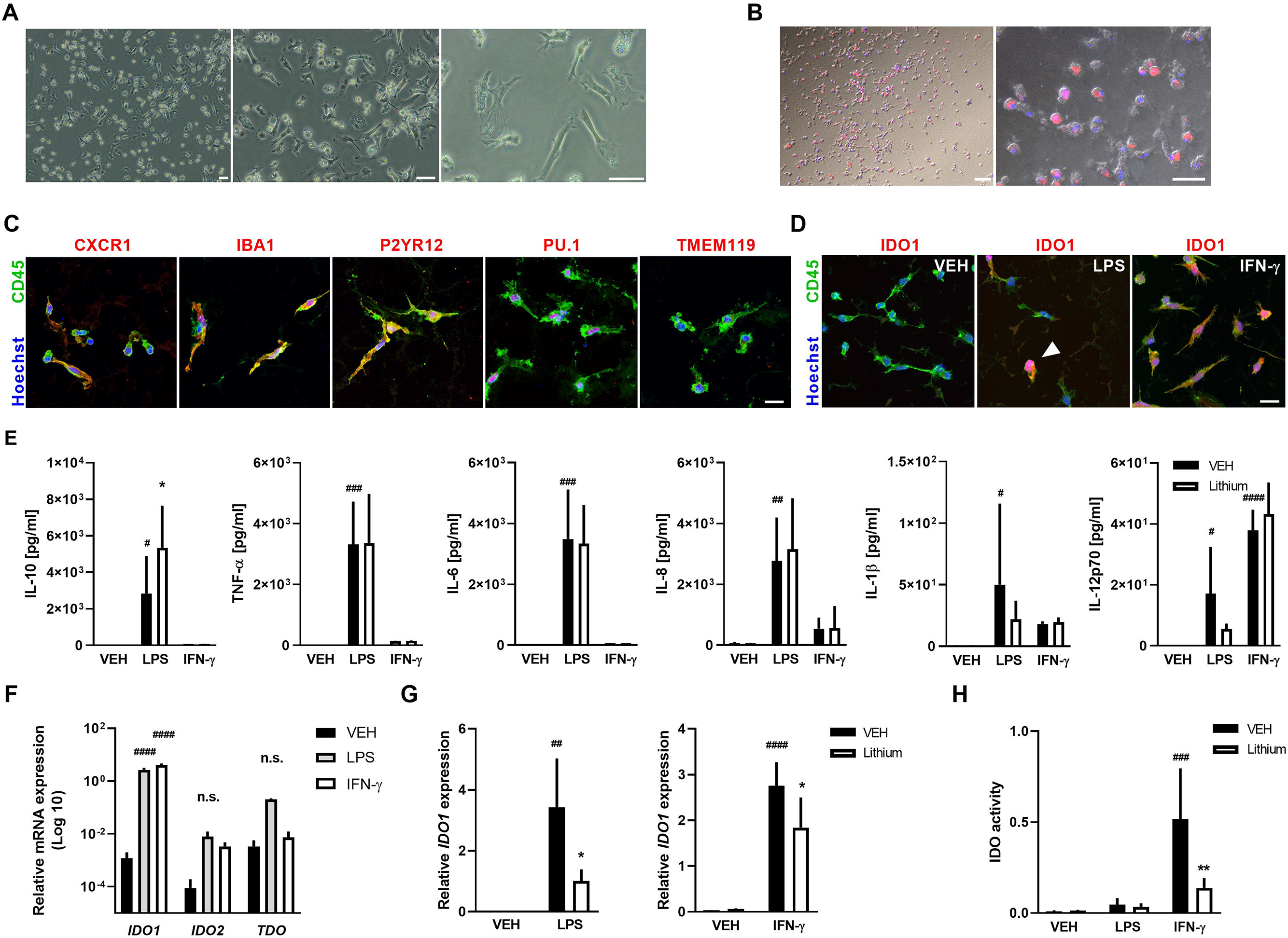
Tryptophan catabolism in hiPSC-derived microglia. **(A)** Representative brightfield images of hiPSC-derived microglia. Scale bar: 50 μm **(B)** Live cell images of hiPSC-derived microglia phagocytosing bacterial particles (red), blue: nuclear stain Hoechst 33342. Brightfield and fluorescence images were merged into a composite image. Scale bar left: 100 μm, right: 50 μm **(C)** Representative confocal images of differentiated hiPSC-derived microglia-like cells demonstrate expression of typical microglia markers such as CD45, CXCR1, P2YR12, IBA1 and PU.1, but not TMEM119. Hoechst 33342 was used as nuclear counterstain. Scale bar: 20 μm. **(D)** IDO1 protein expression is strongly induced in IFN-γ stimulated cells. Scale bar: 20 μm **(E)** Microglia-like cells derived from hiPSCs show typical release of pro-inflammatory cytokines after 24 h LPS treatment. HiPSC-derived microglia cells from 3 different donor cell lines were used (i.e., n=3 experiments), two-way ANOVA followed by uncorrected Fisher’s LSD, ^#^p<0.05, ^##^p<0.01, ^###^p<0.001, ^####^p<0.0001 vehicle vs. LPS or IFN-γ, *p<0.05 LPS vehicle vs. LPS and co-treatment with lithium. **(F)***IDO1*, *IDO2*, and *TDO* mRNA expression in differentiated hiPSC-derived microglia-like cells 24 h after LPS or IFN-γ treatment. Results are displayed on a logarithmic scale. Significant regulation was only observed for *IDO1*. N=3 experiments, two-way ANOVA followed by uncorrected Fisher’s LSD, ^####^p<0.0001 vehicle vs. LPS or IFN-γ. **(G)** I*DO1* mRNA following 24 h stimulation with LPS (left) or IFN-γ (right). N=3 experiments, two-way ANOVA followed by uncorrected Fisher’s LSD, ^##^p<0.01 vehicle vs. LPS, ^####^p<0.0001 vehicle vs. IFN-γ, *p<0.05 stimulation condition vs. stimulation condition and co-treatment with lithium. **(H)** IDO activity following 24h LPS or IFN-γ treatment, n=3, two-way ANOVA followed by uncorrected Fisher’s LSD, ^###^p<0.001 vehicle vs. IFN-γ, **p<0.01 IFN-γ vs. IFN-γ and co-treatment with lithium.

## Discussion

The biological mechanisms underpinning lithium therapy are not clearly understood. One of the most robust and consistent findings reported in the literature and replicated here is that lithium inhibits glycogen synthase kinase (GSK)3β (Chalecka-Franaszek and Chuang, 1999; Klein and Melton, 1996; Stambolic et al., 1996), a ubiquitous serine/threonine kinase with particularly high expression in brain (Woodgett, 1990). A number of clinical studies have demonstrated an increase in 5-hydroxyindoleacetic acid (5-HIAA), the main serotonin metabolite, in cerebrospinal fluid (CSF) of patients receiving lithium (Berrettini et al., 1985; Bowers and Heninger, 1977; Fyro et al., 1975). Co-prescription of lithium and serotonergic antidepressants heightens the risk of serotonin toxicity (Adan-Manes et al., 2006; Prakash et al., 2017; Sobanski et al., 1997). Furthermore, accumulating evidence has linked higher-lethality suicide attempts to central serotonin system hypofunction (e.g. (Asberg et al., 1976; Mann et al., 1989; Roy et al., 1989; Sullivan et al., 2015)) whereas lithium has proven anti-suicidal properties (Cipriani et al., 2013). Based on these findings, it has long been suspected that part of lithium’s benefit might be mediated directly via serotonin (Muller-Oerlinghausen, 1985). Against this background, our study was designed to investigate in microglia, the brain’s resident immune cells, the effect of lithium on the inflammation-induced kynurenine pathway, which siphons tryptophan away from serotonin biosynthesis.

Microglia are the main effector cell driving the brain’s innate immune response. As such, microglia are an important first line of defense and lend themselves well as a cellular target for strategies aimed at modulating neuroinflammation (Colonna and Butovsky, 2017). To our knowledge, this is the first report documenting the effect of lithium on kynurenine-pathway activity in microglia.

Considerable doubt has been raised as to whether mice are suitable to model inflammation observed in human patients (Seok et al., 2013). Notwithstanding the extra labor involved, the experiments reported herein were therefore conducted in human-derived cells. Cell culture models are inherently reductionist, being composed of only one cell type investigated *ex vivo* under non-physiological conditions for a necessarily relatively short period of time. The attendant difficulties and limitations should be clearly acknowledged. Not infrequently, the pace of such experiments in the dish exceeds by far the speed with which the corresponding effects become observable in vivo. As concerns microglia cultures in particular, in the interest of reasonably quick and reproducible experiments, biomolecules and pharmacological agents are typically applied to cells at rather high concentrations, the equivalents of which would not usually be used in experimental animals or expected to be seen in the human brain unless in most severe disease. This is certainly true for the concentrations of LPS and IFN-γ used herein for a short span of just 24 hours in line with numerous prior studies (Carrillo-Jimenez et al., 2019; Frugier et al., 2010; Huang et al., 2009; Nguyen et al., 2017). It should also be noted that the effects of lithium investigated in our experiments all relate to intracellular signaling and that IDO1, one of whose main evolutionary roles seems to have been to starve intracellular pathogens of tryptophan, is a cytosolic protein lacking a known secreted isoform (Schmidt and Schultze, 2014; van Baren and Van den Eynde, 2015). The intracellular concentration of lithium in microglia of experimental animals or human patients receiving therapeutic lithium salts remains to be established. Moreover, it should not go unmentioned that a recent study using ^7^Li MR imaging found significant heterogeneity in lithium distribution within the brain of subjects with bipolar disorder with particularly high signal intensity found, most remarkably, in the brain stem (Smith et al., 2018). Further, ^7^Li MR spectroscopy suggests that an extensive fraction of lithium resides in the intracellular compartment in the human brain in vivo, albeit with estimates falling within a broad range (Komoroski et al., 1997). Finally, a steady-state concentration of lithium (i.e., an intracellular concentration producing stable therapeutic effects) is only achieved in vivo after four to five days (Ward et al., 1994). Against this complex backdrop, we were guided in our choice of lithium concentration primarily by the previous literature on lithium effects in microglia and glial cells in culture and proceeded after having confirmed good tolerability using the MTT assay (e.g. (Cao et al., 2017; Davenport et al., 2010; Feinstein, 1998; Yuskaitis and Jope, 2009; Zheng et al., 2017)).

Our study establishes a signaling pathway by which lithium inhibits inflammatory activation of the kynurenine pathway in human microglia (Figure 5). The effects of lithium were corroborated using multiple readouts including mRNA and protein expression as well as the KYN/TRP ratio.

**Figure 5.**
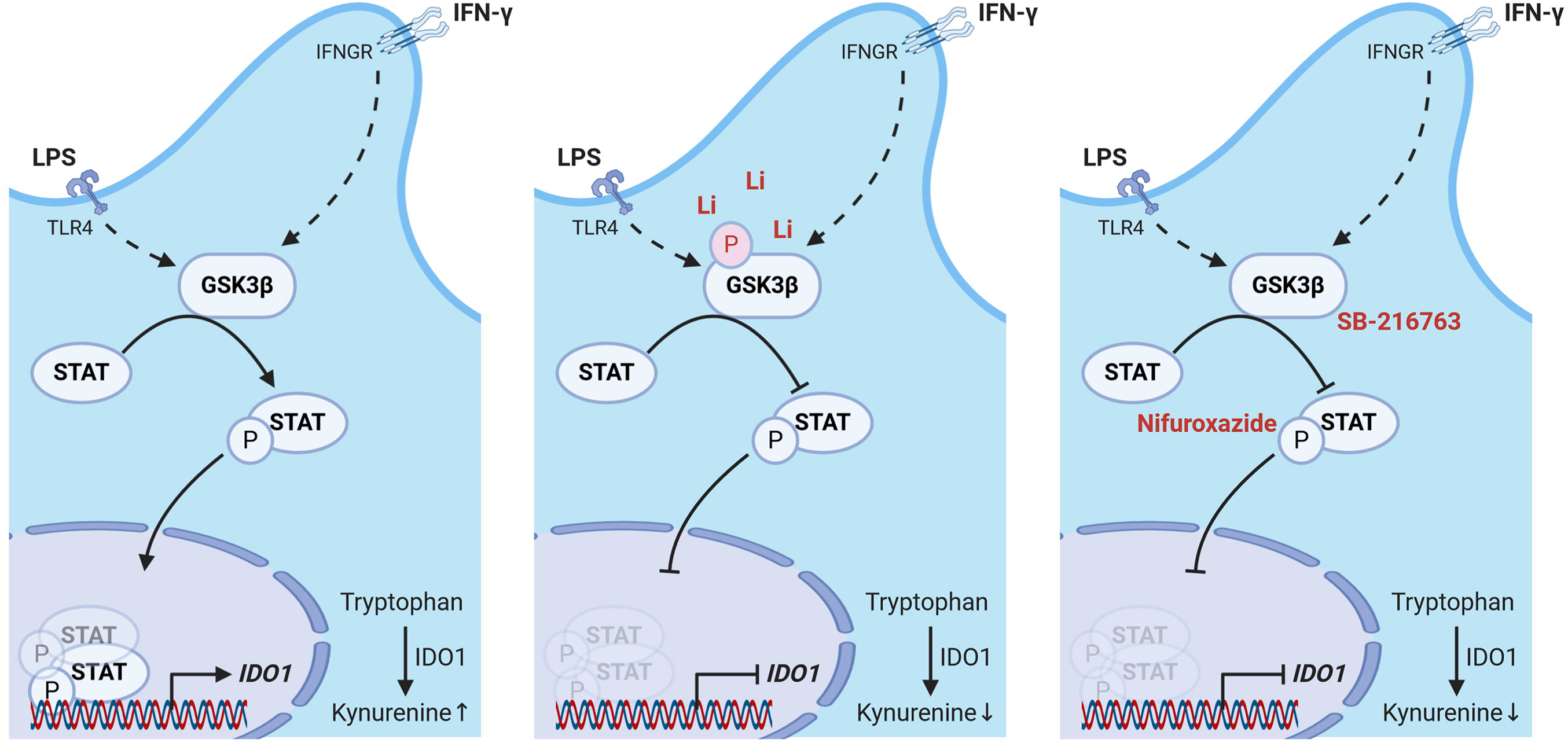
Lithium inhibits tryptophan catabolism along the kynurenine pathway in activated human microglia. When microglia become activated in response to stimulation of toll-like receptor 4 (TLR4) or the IFN-γ receptor (IFNGR), GSK3β phosphorylates STAT1 at serine 727 and STAT3 at tyrosine 705, allowing dimerization, nuclear translocation, and inflammation-induced *IDO1* mRNA transcription. Lithium phosphorylates GSK3β at serine 9, thereby blocking the crucial role of GSK3β in microglia activation. GSK3β inhibitor SB-216763 and STAT inhibitor nifuroxazide recapitulate the effects of lithium on *IDO1* mRNA transcription. Created with BioRender.com

Glycogen synthase kinase 3β (GSK3β) plays a central role in regulating the cellular immune response. In line with the existing literature, we confirmed inhibitory phosphorylation of GSK3β at serine 9 in human microglia treated with lithium (Stambolic et al., 1996; Sutherland et al., 1993). In mice, GSK3β activation after stimulation with IFN-γ has been shown to lead to downstream phosphorylation of STAT1 at serine 727 (Tsai et al., 2009) and STAT3 at tyrosine 705 (Beurel and Jope, 2008). Using an antibody array to interrogate our samples, we confirmed this inflammatory signaling cascade in human cells. Importantly, enriched binding of both STAT1 and STAT3 to the *IDO1* promoter was observed after IFN-γ activation of immortalized human microglia cells. Further, we identified dephosphorylation of STAT1 at serine 727 and STAT3 at tyrosine 705 as the principal signaling route of lithium, blocking nuclear translocation and DNA binding of these transcription factors in response to inflammatory activation (Darnell et al., 1994). Crucially, we were able to show that pharmacological inhibition of GSK3 and STAT recapitulates lithium’s inhibitory effect on the kynurenine pathway.

Taken together, immortalized human microglia, primary human microglia harvested from surgical specimens, and microglia-like cells differentiated from human iPSCs lend themselves well to modeling key features of neuroinflammation. The main difference between the three cell types relates to stimulation with LPS, which produced marked induction of IDO1 in primary microglia, showed some upregulation in hiPSC-derived cells, but yielded virtually no response in the immortalized human microglia cell line investigated. *IDO1* mRNA and protein expression as well as IDO activity were robustly induced after stimulation with IFN-γ. Our study suggests that, in time, hiPSC-derived microglia might become a useful tool that will contribute to personalized medicine. Uses might include, for example, patient-specific characterization of microglia responses to inflammatory stimuli with a view to predicting psychiatric side effects of immunomodulatory therapies such as interferon-induced depression.

In sum, lithium counteracts tryptophan catabolism along the kynurenine pathway in stimulated human-derived microglia by increasing inhibitory GSK3β^S9^ phosphorylation and reducing, in turn, STAT1^S727^ and STAT3^Y705^ phosphorylation levels. Although lithium is a highly pleiotropic agent that interacts with multiple intracellular signaling pathways, we speculate that inhibition of inflammation-induced tryptophan breakdown explains critical aspects of lithium’s clinical and pharmacological properties.

## Acknowledgments

We thank Stefanie Balz, Bettina Herrmann, and Renate Franke for excellent technical assistance. We are indebted to Frank Heppner and Marina Jendrach for allowing us to use the Mesoscale reader. We thank the Berlin Institute of Health stem cell core facility (Harald Stachelscheid, Judit Küchler, Kristin Fischer) for provision of hiPSC lines and support.

This work was supported by the Deutsche Forschungsgemeinschaft (EXC 2049-390688087 to M.E.; KR 2956/4-1 and KR 2956/6-1 to G.K.; GE 2576/3-1 and GE 2576/5-1 to K.G.), the Bundesministerium für Bildung und Forschung (CSB to M.E., K.G. and G.K.), the German Center for Neurodegenerative Diseases (DZNE to M.E.), the German Centre for Cardiovascular Research (DZHK to M.E. and K.G.), the van Geest Cardiovascular Development Fund (to G.K.), and the Corona Foundation (to M.E.).

## Methods

Key Resources Table (Table S3)

### Resource availability

#### Lead Contact

Further information and requests for resources and reagents should be directed to and will be fulfilled by the Lead Contact, Karen Gertz.

#### Materials Availability

This study did not generate new unique reagents.

#### Data and Code Availability

This study did not generate any unique datasets or code.

### Experimental Model and Subject Details

#### Human brain specimens

Human brain cortical biopsy material was obtained, after prior written informed consent, from four patients undergoing respective surgery for medication-resistant epilepsy. All procedures were approved by the local ethics committee of the Charité-Universitätsmedizin Berlin (EA2/111/14) and performed in accordance with the Declaration of Helsinki.

#### Immortalized Human Microglia

Immortalized human microglia (HM-IM; Innoprot) were cultivated on collagen A coated plasticware in microglia medium (Innoprot) for up to 8 passages. Cells were seeded and cultured at a density of 3 × 10^4^ cells / cm^2^ until they reached a confluence of 90%. To stop proliferation, cells were incubated for an additional 24 h in RPMI 1640 medium (Biochrom, Merck) lacking fetal calf serum before further experimentation.

#### Human induced pluripotent stem cells (hiPSCs)

HiPSC lines (BIHi005-A, BIHi004-A, and BIHi001-A, see hPSCreg database for details https://hpscreg.eu/) were provided by the Berlin Institute of Health (BIH) stem cell core facility. HiPSCs were maintained in 6-well plates (Corning) in feeder-free conditions using growth factor-reduced Geltrex (Thermo Fisher Scientific) coated culture dishes in complete TeSR-E8 medium (Stem Cell Technologies) and cultured in a humidified incubator (5% CO_2_, 37°C). HiPSCs were fed daily with fresh media and passaged every 7–8 days.

### Method Details

#### Reagents

Plasticware was coated with with poly-L-lysin (diluted 1:20 in distilled water; Biochrom, Merck) or collagen A (0.5 mg/ml Biochrom, Merck) overnight at 4°C followed by washes with 1xDPBS (Gibco™, Thermo Fisher Scientific). Cells were activated with LPS (1 μg/ml; Sigma-Aldrich) or human IFN-γ (200 ng/ml; Peprotech). Lithium chloride (10 mM; Sigma-Aldrich), SB-216763 (10 μM; AdipoGen), and nifuroxazide (10 μM, Sigma-Aldrich) were applied 30 min before activation.

#### Primary postnatal human microglia cultures

For the isolation of human microglia from surgical brain specimens, we essentially followed a previously published protocol (Rustenhoven et al., 2016). Briefly, blood vessels and meninges were removed from resected brain tissue using a dissection microscope (Zeiss Stemi DV4). Tissue was weighed, diced and dissociated using the Neural Tissue Dissociation Kit (P) (#130-092-628, Miltenyi Biotec). Microglia were cultured as adherent cells in poly-L-lysin coated T75 tissue culture flasks in microglia culture medium containing DMEM/F12 (Gibco™, Thermo Fisher Scientific), 10% fetal calf serum (Biochrom, Merck) and 1X penicillin-streptomycin solution (Biochrom, Merck). Purity of cultures was verified by immunostaining (Figure S2).

#### Human iPSC derived microglia

Differentiation of hiPSCs into hematopoietic progenitor cells (iHPCs) and subsequent differentiation of iHPCs into hiPSC derived microglia was performed as described earlier (Abud et al., 2017). Briefly, hiPSCs were first differentiated into hematopoetic progenitor cells over a time period of 11 days using hematopoetic differentiation medium (HDM) containing IMDM (50%), F12 (50%), ITSG-X (2% v/v), Glutamax (1X), chemically-defined lipid concentrate (1X), non-essential amino acids (NEAA; 1X), and Penicillin/Streptomycin (P/S; 1% V/V; all from Thermo Fisher Scientific), L-ascorbic acid 2-phosphate magnesium (64 μg/ml), monothioglycerol (400 μM), and PVA (10 μg/ml; all from Sigma). HDM was supplemented on day 0 with FGF2 (50 ng/ml; Peprotech), BMP4 (50 ng/ml; Miltenyi Biotec), Activin-A (12.5 ng/ml; Peprotech), RI (1 μm; Stem Cell Technology) and LiCl (2 mM; Sigma-Aldrich). Day 2 HDM was supplemented with FGF2 (50 ng/ml) and VEGF (50 ng/ml), days 4, 6, and 8 with FGF2 (50 ng/ml), VEGF (50 ng/ml), TPO (50 ng/ml), SCF (10 ng/ml), IL-6 (50 ng/ml), and IL-3 (10 ng/ml; all from Peprotech). Day 10 cells were harvested and FACsorted based on CD43 expression. CD43+ cells were transferred into microglia differentiation medium (MDM) containing phenol-free DMEM/F12 (1:1), ITS-G (2%v/v), B27 (2% v/v), N2 (0.5%, v/v; all from Thermo Fisher Scientific), monothioglycerol (200 μM), Glutamax (1X), NEAA (1X), and additional insulin (5 μg/ml; Sigma) for another 28 days. MDM contained the following supplements: Day 10-35: M-CSF (25 ng/ml), IL-34 (100 ng/ml), and TGFb-1 (50 ng/ml; all from Peprotech). Day 35-38: M-CSF (25 ng/ml), IL-34 (100 ng/ml;), TGFb-1 (50 ng/ml), CD200 (100 ng/ml, Novoprotein), and CX3CL1 (100 ng/ml; Peprotech). After 38 days of differentiation, hiPSC derived microglia cells were harvested and seeded in 96 well plates (20,000 cells/well; Corning) containing MDM and M-CSF (25 ng/ml), IL-34 (100 ng/ml), TGFb-1 (50 ng/ml), CX3CL1 (100 ng/ml) and CD200 (100 ng/ml).

#### Immunofluorescence and quantification

Cells were seeded onto poly-L-lysine precoated chamber slides (5000 cells/well; Sarstedt, Nümbrecht, Germany). Briefly, in preparation of staining, cells were washed three times with DPBS (1x) and then fixed with PFA (4% w/v) overnight at 4°C. Primary and secondary antibodies were diluted in TBS blocking buffer (96 ml 1xTBS, 1 ml 10% Triton X-100, 3 ml donkey serum). Primary antibodies were applied in the following concentrations: mouse anti-CD45 (EXBIO) 1:100, rabbit anti-IDO1 (Cell Signaling Technology) 1:100, rabbit anti-IBA1 (Wako Chemicals) 1:500, rabbit anti-PU.1 (Cell Signaling) 1:200, rabbit anti-TMEM119 (Biozol) 1:50, and rabbit anti-P2RY12 (Biozol) 1:50. Secondary antibodies coupled to different fluorochromes (Alexa488, Alexa647, Alexa488) were all from Invitrogen (Thermo Fisher Scientific, Waltham, M.A.). Secondary antibodies were used in a dilution of 1:400.

To measure mean fluorescence intensity of IDO1, Z-stack images were collected on an LSM 700 confocal microscope (Carl Zeiss). Consistent laser power and microscope settings were used across experiments. Zen® software (Carl Zeiss) was used for image analysis.

#### Isolation of mRNA and quantitative polymerase chain reaction

RNA was isolated from cultured cells using the NucleoSpin^®^ RNA XS kit (Macherey-Nagel). M-MLV reverse transcriptase and random hexamers (both from Promega, Thermo Fisher Scientific) were used for reverse transcription of RNA into cDNA. For polymerase chain reaction amplification, we used gene-specific primers and Light Cycler^®^ 480 SYBR Green I Master (Roche Diagnostics). Polymerase chain reaction conditions were as follows: preincubation 95°C, 10 min; 95°C, 10 s, primer-specific annealing temperature, 10 s, 72°C, 15 s (45 cycles). Crossing points of amplified products were determined using the Second Derivative Maximum Method (Light Cycler Version LCS480 1.5.0.39, Roche). Quantification of messenger RNA expression was relative to receptor expression-enhancing protein 5 (*REEP5*; (Eisenberg and Levanon, 2013)). The specificity of polymerase chain reaction products was checked using melting curve analysis. PCR products were run on a 1.5% agarose gel to demonstrate the presence of a single amplicon of the correct size. Furthermore, negative controls (i.e., reaction mix lacking either template DNA or reverse transcriptase) yielded no bands on the gel.

#### Western Blot analysis

For protein extraction, cells were harvested in RIPA lysis buffer (150 mM NaCl, 5 mM EDTA, 1% NP-40, 1% sodium deoxycholate, 0.1% SDS, 25 mM Tris-HCl pH 7.6) containing phosphatase and protease inhibitors (PhosSTOP™, cOmplete™; both from Roche). Protein concentration was determined with the Pierce™ BCA Protein Assay Kit (Pierce™, Thermo Fisher Scientific). Equal amounts of protein were loaded on 10% SDS-polyacrylamide gels (Bio-Rad Laboratories, Hercules, CA) and blotted onto PVDF membranes (Immobilon, Millipore, Merck) for 150 min at 20 V. Blots were probed with primary and secondary antibodies and developed using near-infrared (NIR) fluorescence detection with Odyssey^®^ CLx Imaging System (LI-COR). Equal loading of protein was confirmed by blotting against GAPDH or ACTB. Primary and secondary antibodies were applied in the following dilutions: IDO1 (1 mg/ml;) 1:1,000, GSK3β (Cell Signaling) 1:250, Phospho-GSK3β (Ser9; Cell Signaling) 1:250, STAT1 (Cell Signaling) 1:500, Phospho-STAT1 (Ser727; AAT Bioquest) 1:500, STAT3 (Cell Signaling) 1:500; Phospho-STAT3 (Tyr705; Cell Signaling) 1:500; GAPDH (Cell Signaling) 1:1,000; ACTB (Abcam) 1:1,000; IRDye^®^ 800 (LI-COR) 1:10,000 and IRDye^®^ 680 (LI-COR) 1:10,000.

#### Intracellular immune cell signaling array

Cells were harvested in lysis buffer (Cell Signaling) and stored at −80°C until analysis. The PathScan®Immune antibody array kit (Cell Signaling) was used according to the manufacturer’s instructions. Fluorescent images were acquired using the LI-COR® Biosciences Odyssey® imaging system. Fluorescence intensities for each spot were quantified using the LI-COR® Image Studio array analysis software and depicted as bar graphs (LI-COR® Biosciences).

#### Phagocytosis Assay

To observe phagocytosis of bacterial particles, microglia cells cultured on PLL coated plasticware were incubated in live cell imaging solution (Thermo Fisher Scientific) with pHrodo Red S. aureus (Thermo Fisher Scientific) according to manufacturer′s manual. After 2 h incubation at 37°C, Hoechst dye 33342 (2 μm; Abcam) was added for 10 min at 37°C, washed once with live cell imaging solution and imaged using an inverted fluorescence microscope (DMI 3000, Leica).

#### MTT Assay

Cell viability was assayed after 24 h incubation with LPS or IFN-γ as previously described (Uhlemann et al., 2016). The MTT labelling agent (Sigma-Aldrich) was added to the cells at a final concentration of 0.5 mg/ml. The converted dye was solubilized in 10 %SDS in 0.01 M HCl and measured at 550 nm with a platereader (TriStar LB941, Berthold Technologies).

#### Cytokine measurements

Cytokines in culture medium were measured using the Mesoscale MSD^©^ Multi-Spot Assay System and the SECTOR Imager 6000 plate reader (both from Meso Scale Diagnostics) according to the manufacturer’s instructions. All samples were run at least in duplicate.

#### Chromatin immunoprecipitation (ChIP)

Chromatin for ChIP was prepared from immortalized human microglia using the Imprint® Chromatin Immunoprecipitation kit (Sigma-Aldrich) according to the manufacturer’s instructions. Briefly, fragmented chromatin was immunoprecipitated with a ChIP-grade antibody against either STAT1 or STAT3 (both from BioLegend). Following reversal of cross-links and DNA precipitation, enriched DNA was analyzed by PCR amplification using a region-specific ChIP primer. QPCR data are presented as percent of input (%IP).

#### Quantification of tryptophan and kynurenine concentrations

Cell culture medium (50 μl) was diluted three times with 0.3 M HClO_4_ and centrifuged for 15 min at 13,000 × *g* and 4 °C. Supernatant was transferred to a Spin-X^®^ centrifuge tube filter (0.22 μm; Corning) and centrifuged at 5,000 × *g* for 5 min at 4 °C. The flow-through was injected into the HPLC system. Kynurenine and tryptophan were analyzed by isocratic high-performance liquid chromatography (HPLC) using UltiMate 3000 pump and autosampler connected to a guard cell 5020 (+750 mV) and Coulochem III electrochemical detector with high sensitivity analytical cell 5011A (ECD1, +450 mV; ECD2, +700 mV; all from Thermo Fisher Scientific, Inc.). Both analytes were separated on a Hypersil™ BDS C18 column (150 mm, VDS-optilab) using isocratic flow of the mobile phase MD-TM (Thermo Fisher Scientific) at a rate of 0.5 ml/min. Each run included standard curves of L-kynurenine and L-tryptophan (Sigma-Aldrich; diluted in 0.1 M HCl) with concentrations ranging from 10-500 ng/ml. For data acquisition and analysis, Chromeleon™ 7.2 Chromatography Data System (CDS) software was used (Thermo Fisher Scientific).

### Quantification and Statistical Analysis

#### Statistical analysis

Values are presented as mean ± SD. All statistical analyses were performed using GraphPad Prism 8 (GraphPad Software). Shapiro-Wilk test was used to confirm normal distribution of data. Unpaired t-test or two-way ANOVA were used to determine statistical significance. The results of relevant post hoc tests are given in the text. Percent effect was calculated relative to vehicle-treated controls (100%). P<0.05 was considered as statistically significant.

## References

Abud, E.M., Ramirez, R.N., Martinez, E.S., Healy, L.M., Nguyen, C.H.H., Newman, S.A., Yeromin, A.V., Scarfone, V.M., Marsh, S.E., Fimbres, C., et al. (2017). iPSC-Derived Human Microglia-like Cells to Study Neurological Diseases. Neuron 94, 278–293 e279.

Adan-Manes, J., Novalbos, J., Lopez-Rodriguez, R., Ayuso-Mateos, J.L., and Abad-Santos, F. (2006). Lithium and venlafaxine interaction: a case of serotonin syndrome. J Clin Pharm Ther 31, 397–400.

Ajmone-Cat, M.A., D’Urso, M.C., di Blasio, G., Brignone, M.S., De Simone, R., and Minghetti, L. (2016). Glycogen synthase kinase 3 is part of the molecular machinery regulating the adaptive response to LPS stimulation in microglial cells. Brain Behav Immun 55, 225–235.

Asberg, M., Traskman, L., and Thoren, P. (1976). 5-HIAA in the cerebrospinal fluid. A biochemical suicide predictor? Arch Gen Psychiatry 33, 1193–1197.

Bauer, M., Adli, M., Bschor, T., Pilhatsch, M., Pfennig, A., Sasse, J., Schmid, R., and Lewitzka, U. (2010). Lithium’s emerging role in the treatment of refractory major depressive episodes: augmentation of antidepressants. Neuropsychobiology 62, 36–42.

Berrettini, W.H., Nurnberger, J.I., Jr., Scheinin, M., Seppala, T., Linnoila, M., Narrow, W., Simmons-Alling, S., and Gershon, E.S. (1985). Cerebrospinal fluid and plasma monoamines and their metabolites in euthymic bipolar patients. Biol Psychiatry 20, 257–269.

Beurel, E., and Jope, R.S. (2008). Differential regulation of STAT family members by glycogen synthase kinase-3. J Biol Chem 283, 21934–21944.

Bowers, M.B., Jr., and Heninger, G.R. (1977). Lithium: clinical effects and cerebrospinal fluid acid monoamine metabolites. Commun Psychopharmacol 1, 135–145.

Bronstein, R., Torres, L., Nissen, J.C., and Tsirka, S.E. (2013). Culturing microglia from the neonatal and adult central nervous system. J Vis Exp, 50647.

Cao, Q., Karthikeyan, A., Dheen, S.T., Kaur, C., and Ling, E.A. (2017). Production of proinflammatory mediators in activated microglia is synergistically regulated by Notch-1, glycogen synthase kinase (GSK-3beta) and NF-kappaB/p65 signalling. PLoS One 12, e0186764.

Carrillo-Jimenez, A., Deniz, O., Niklison-Chirou, M.V., Ruiz, R., Bezerra-Salomao, K., Stratoulias, V., Amouroux, R., Yip, P.K., Vilalta, A., Cheray, M., et al. (2019). TET2 Regulates the Neuroinflammatory Response in Microglia. Cell Rep 29, 697–713 e698.

Chalecka-Franaszek, E., and Chuang, D.M. (1999). Lithium activates the serine/threonine kinase Akt-1 and suppresses glutamate-induced inhibition of Akt-1 activity in neurons. Proceedings of the National Academy of Sciences of the United States of America 96, 8745–8750.

Cipriani, A., Hawton, K., Stockton, S., and Geddes, J.R. (2013). Lithium in the prevention of suicide in mood disorders: updated systematic review and meta-analysis. BMJ 346, f3646.

Coghlan, M.P., Culbert, A.A., Cross, D.A., Corcoran, S.L., Yates, J.W., Pearce, N.J., Rausch, O.L., Murphy, G.J., Carter, P.S., Roxbee Cox, L., et al. (2000). Selective small molecule inhibitors of glycogen synthase kinase-3 modulate glycogen metabolism and gene transcription. Chem Biol 7, 793–803.

Colonna, M., and Butovsky, O. (2017). Microglia Function in the Central Nervous System During Health and Neurodegeneration. Annu Rev Immunol 35, 441–468.

Darnell, J.E., Jr., Kerr, I.M., and Stark, G.R. (1994). Jak-STAT pathways and transcriptional activation in response to IFNs and other extracellular signaling proteins. Science 264, 1415–1421.

Davenport, C.M., Sevastou, I.G., Hooper, C., and Pocock, J.M. (2010). Inhibiting p53 pathways in microglia attenuates microglial-evoked neurotoxicity following exposure to Alzheimer peptides. J Neurochem 112, 552–563.

Dong, H., Zhang, X., Dai, X., Lu, S., Gui, B., Jin, W., Zhang, S., Zhang, S., and Qian, Y. (2014). Lithium ameliorates lipopolysaccharide-induced microglial activation via inhibition of toll-like receptor 4 expression by activating the PI3K/Akt/FoxO1 pathway. Journal of neuroinflammation 11, 140.

Eisenberg, E., and Levanon, E.Y. (2013). Human housekeeping genes, revisited. Trends Genet 29, 569–574.

Feinstein, D.L. (1998). Potentiation of astroglial nitric oxide synthase type-2 expression by lithium chloride. J Neurochem 71, 883–886.

Frugier, T., Morganti-Kossmann, M.C., O’Reilly, D., and McLean, C.A. (2010). In situ detection of inflammatory mediators in post mortem human brain tissue after traumatic injury. J Neurotrauma 27, 497–507.

Fyro, B., Petterson, U., and Sedvall, G. (1975). The effect of lithium treatment on manic symptoms and levels of monoamine metabolites in cerebrospinal fluid of manic depressive patients. Psychopharmacologia 44, 99–103.

Geddes, J.R., Burgess, S., Hawton, K., Jamison, K., and Goodwin, G.M. (2004). Long-term lithium therapy for bipolar disorder: systematic review and meta-analysis of randomized controlled trials. Am J Psychiatry 161, 217–222.

Huang, W.C., Lin, Y.S., Wang, C.Y., Tsai, C.C., Tseng, H.C., Chen, C.L., Lu, P.J., Chen, P.S., Qian, L., Hong, J.S., et al. (2009). Glycogen synthase kinase-3 negatively regulates anti-inflammatory interleukin-10 for lipopolysaccharide-induced iNOS/NO biosynthesis and RANTES production in microglial cells. Immunology 128, e275–286.

Jeon, S.W., and Kim, Y.K. (2016). Neuroinflammation and cytokine abnormality in major depression: Cause or consequence in that illness? World J Psychiatry 6, 283–293.

Jope, R.S., and Johnson, G.V. (2004). The glamour and gloom of glycogen synthase kinase-3. Trends Biochem Sci 29, 95–102.

Klein, P.S., and Melton, D.A. (1996). A molecular mechanism for the effect of lithium on development. Proceedings of the National Academy of Sciences of the United States of America 93, 8455–8459.

Komoroski, R.A., Pearce, J.M., and Newton, J.E. (1997). The distribution of lithium in rat brain and muscle in vivo by 7Li NMR imaging. Magn Reson Med 38, 275–278.

Kopschina Feltes, P., Doorduin, J., Klein, H.C., Juarez-Orozco, L.E., Dierckx, R.A., Moriguchi-Jeckel, C.M., and de Vries, E.F. (2017). Anti-inflammatory treatment for major depressive disorder: implications for patients with an elevated immune profile and non-responders to standard antidepressant therapy. J Psychopharmacol 31, 1149–1165.

Malhi, G.S., Gessler, D., and Outhred, T. (2017). The use of lithium for the treatment of bipolar disorder: Recommendations from clinical practice guidelines. J Affect Disord 217, 266–280.

Mann, J.J., Arango, V., Marzuk, P.M., Theccanat, S., and Reis, D.J. (1989). Evidence for the 5-HT hypothesis of suicide. A review of post-mortem studies. Br J Psychiatry Suppl, 7–14.

McKim, D.B., Weber, M.D., Niraula, A., Sawicki, C.M., Liu, X., Jarrett, B.L., Ramirez-Chan, K., Wang, Y., Roeth, R.M., Sucaldito, A.D., et al. (2018). Microglial recruitment of IL-1beta-producing monocytes to brain endothelium causes stress-induced anxiety. Mol Psychiatry 23, 1421–1431.

Muller-Oerlinghausen, B. (1985). Lithium long-term treatment--does it act via serotonin? Pharmacopsychiatry 18, 214–217.

Nahman, S., Belmaker, R.H., and Azab, A.N. (2012). Effects of lithium on lipopolysaccharide-induced inflammation in rat primary glia cells. Innate Immun 18, 447–458.

Nelson, E.A., Walker, S.R., Kepich, A., Gashin, L.B., Hideshima, T., Ikeda, H., Chauhan, D., Anderson, K.C., and Frank, D.A. (2008). Nifuroxazide inhibits survival of multiple myeloma cells by directly inhibiting STAT3. Blood 112, 5095–5102.

Nguyen, H.M., Grossinger, E.M., Horiuchi, M., Davis, K.W., Jin, L.W., Maezawa, I., and Wulff, H. (2017). Differential Kv1.3, KCa3.1, and Kir2.1 expression in “classically” and “alternatively” activated microglia. Glia 65, 106–121.

Nie, X., Kitaoka, S., Tanaka, K., Segi-Nishida, E., Imoto, Y., Ogawa, A., Nakano, F., Tomohiro, A., Nakayama, K., Taniguchi, M., et al. (2018). The Innate Immune Receptors TLR2/4 Mediate Repeated Social Defeat Stress-Induced Social Avoidance through Prefrontal Microglial Activation. Neuron 99, 464–479 e467.

Perez-Cruet, J., Tagliamonte, A., Tagliamonte, P., and Gessa, G.L. (1971). Stimulation of serotonin synthesis by lithium. J Pharmacol Exp Ther 178, 325–330.

Prakash, S., Adroja, B., and Parekh, H. (2017). Serotonin syndrome in patients with headache disorders. BMJ Case Rep 2017.

Roy, A., De Jong, J., and Linnoila, M. (1989). Cerebrospinal fluid monoamine metabolites and suicidal behavior in depressed patients. A 5-year follow-up study. Arch Gen Psychiatry 46, 609–612.

Rustenhoven, J., Park, T.I., Schweder, P., Scotter, J., Correia, J., Smith, A.M., Gibbons, H.M., Oldfield, R.L., Bergin, P.S., Mee, E.W., et al. (2016). Isolation of highly enriched primary human microglia for functional studies. Sci Rep 6, 19371.

Schmidt, S.V., and Schultze, J.L. (2014). New Insights into IDO Biology in Bacterial and Viral Infections. Front Immunol 5, 384.

Seok, J., Warren, H.S., Cuenca, A.G., Mindrinos, M.N., Baker, H.V., Xu, W., Richards, D.R., McDonald-Smith, G.P., Gao, H., Hennessy, L., et al. (2013). Genomic responses in mouse models poorly mimic human inflammatory diseases. Proc Natl Acad Sci U S A 110, 3507–3512.

Smith, F.E., Thelwall, P.E., Necus, J., Flowers, C.J., Blamire, A.M., and Cousins, D.A. (2018). 3D (7)Li magnetic resonance imaging of brain lithium distribution in bipolar disorder. Mol Psychiatry 23, 2184–2191.

Smith, K.A., and Cipriani, A. (2017). Lithium and suicide in mood disorders: Updated meta-review of the scientific literature. Bipolar Disord 19, 575–586.

Sobanski, T., Bagli, M., Laux, G., and Rao, M.L. (1997). Serotonin syndrome after lithium add-on medication to paroxetine. Pharmacopsychiatry 30, 106–107.

Stambolic, V., Ruel, L., and Woodgett, J.R. (1996). Lithium inhibits glycogen synthase kinase-3 activity and mimics wingless signalling in intact cells. Curr Biol 6, 1664–1668.

Sullivan, G.M., Oquendo, M.A., Milak, M., Miller, J.M., Burke, A., Ogden, R.T., Parsey, R.V., and Mann, J.J. (2015). Positron emission tomography quantification of serotonin(1A) receptor binding in suicide attempters with major depressive disorder. JAMA Psychiatry 72, 169–178.

Sutherland, C., Leighton, I.A., and Cohen, P. (1993). Inactivation of glycogen synthase kinase-3 beta by phosphorylation: new kinase connections in insulin and growth-factor signalling. Biochem J 296 (Pt 1), 15–19.

Tam, W.Y., and Ma, C.H. (2014). Bipolar/rod-shaped microglia are proliferating microglia with distinct M1/M2 phenotypes. Sci Rep 4, 7279.

Tiihonen, J. (2017). Use of lithium in patients with unipolar depression - Author’s reply. Lancet Psychiatry 4, 663.

Torres-Platas, S.G., Cruceanu, C., Chen, G.G., Turecki, G., and Mechawar, N. (2014). Evidence for increased microglial priming and macrophage recruitment in the dorsal anterior cingulate white matter of depressed suicides. Brain Behav Immun 42, 50–59.

Tsai, C.C., Kai, J.I., Huang, W.C., Wang, C.Y., Wang, Y., Chen, C.L., Fang, Y.T., Lin, Y.S., Anderson, R., Chen, S.H., et al. (2009). Glycogen synthase kinase-3beta facilitates IFN-gamma-induced STAT1 activation by regulating Src homology-2 domain-containing phosphatase 2. J Immunol 183, 856–864.

Uhlemann, R., Gertz, K., Boehmerle, W., Schwarz, T., Nolte, C., Freyer, D., Kettenmann, H., Endres, M., and Kronenberg, G. (2016). Actin dynamics shape microglia effector functions. Brain Struct Funct 221, 2717–2734.

van Baren, N., and Van den Eynde, B.J. (2015). Tryptophan-degrading enzymes in tumoral immune resistance. Front Immunol 6, 34.

Ward, M.E., Musa, M.N., and Bailey, L. (1994). Clinical pharmacokinetics of lithium. J Clin Pharmacol 34, 280–285.

Woodgett, J.R. (1990). Molecular cloning and expression of glycogen synthase kinase-3/factor The EMBO journal 9, 2431–2438.

Yirmiya, R., Rimmerman, N., and Reshef, R. (2015). Depression as a microglial disease. Trends in neurosciences 38, 637–658.

Yuskaitis, C.J., and Jope, R.S. (2009). Glycogen synthase kinase-3 regulates microglial migration, inflammation, and inflammation-induced neurotoxicity. Cell Signal 21, 264–273.

Zheng, H., Jia, L., Liu, C.C., Rong, Z., Zhong, L., Yang, L., Chen, X.F., Fryer, J.D., Wang, X., Zhang, Y.W., et al. (2017). TREM2 Promotes Microglial Survival by Activating Wnt/beta-Catenin Pathway. J Neurosci 37, 1772–1784.

