## Supplementary Figure Legends for "Lithium inhibits tryptophan catabolism via the inflammation-induced kynurenine pathway in human microglia"

**Supplementary Information**

**Supplementary Figure Legends**

**Supplementary Figure 1. Messenger RNA expression of tryptophan/ N′-formylkynurenine converting enzymes and cell viability after lithium treatment.**

**(A, B)** Immortalized human microglia were treated with IFN-γ for 24 h **(A)** While *IDO1* mRNA was robustly induced, *IDO2* and *TDO* did not show significant changes. **(B)** MTT assay confirmed that neither stimulation with IFN-γ nor lithium treatment (10 mM) affected cell viability. **(C, D)** Primary human microglia were stimulated with either LPS or IFN-γ for 24h **(C)** While *IDO1* mRNA was induced, *IDO2* and *TDO* did now show significant changes. **(D)** MTT assay confirmed that neither stimulation (LPS/IFN-γ) nor treatment with lithium affected viability of primary human microglia in culture. N=3 experiments, two-way ANOVA followed by Tukey’s post hoc test. *P<0.05, ****p<0.0001 between vehicle and stimulation condition (i.e., treatment with either LPS or IFN-γ).

**Supplementary Figure 2. Characterization of primary human microglia.**

**(A)** Primary human microglia express microglia markers IBA1, PU.1, TMEM119, P2YR12, and CD45. **(B)** Composite image showing primary human microglia (brightfield) phagocytosing bacterial particles conjugated to pHrodo dye (red). The Hoechst 33342 nuclear counterstain is blue. **(C)** Primary human microglia were cultured on PLL coated plasticware. The images were taken 24 h after treatment with LPS, IFN-γ or vehicle.
