## Supplementary figures and images for "Lithium inhibits tryptophan catabolism via the inflammation-induced kynurenine pathway in human microglia"

### Figure S1

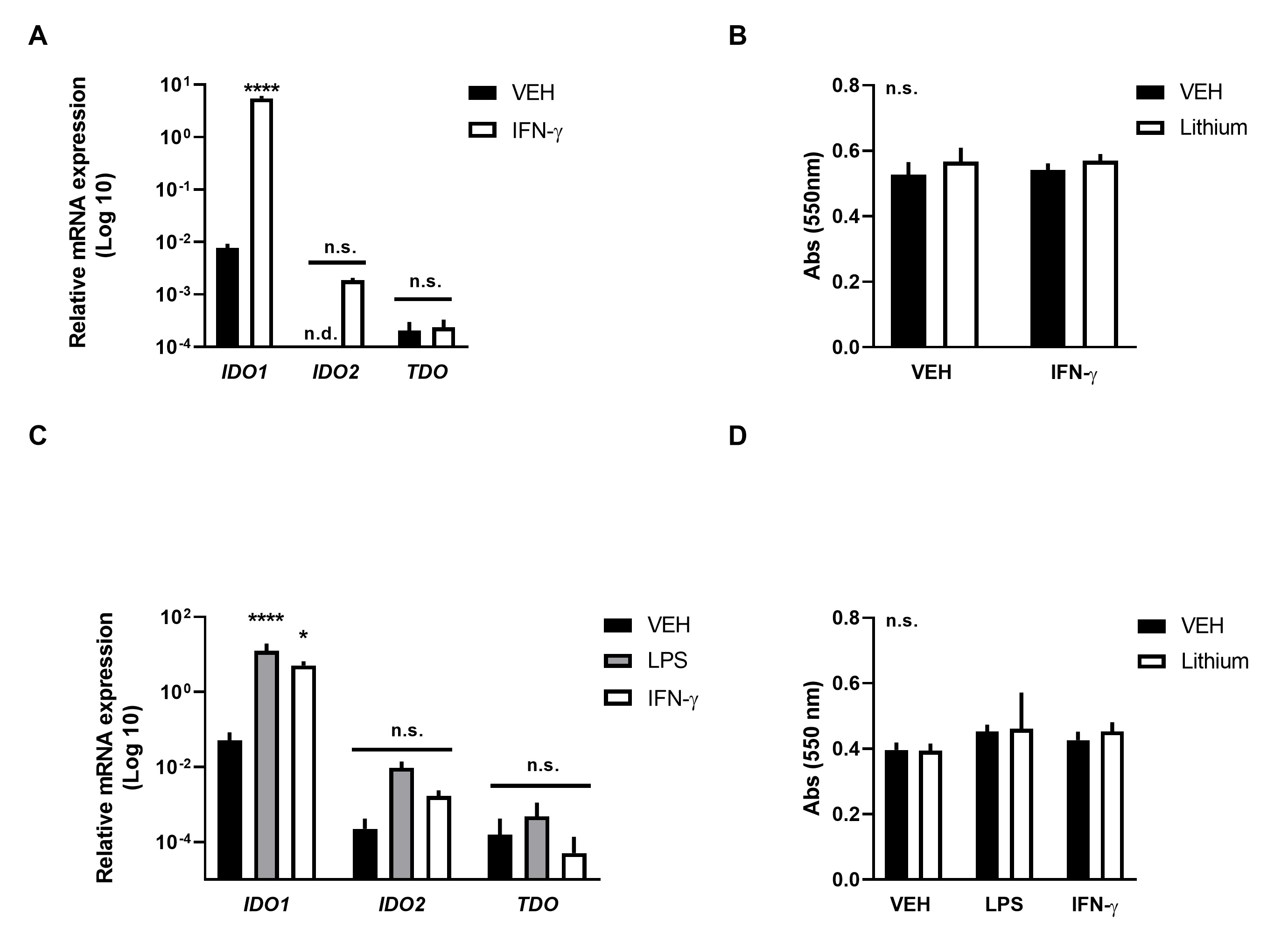

### Figure S2

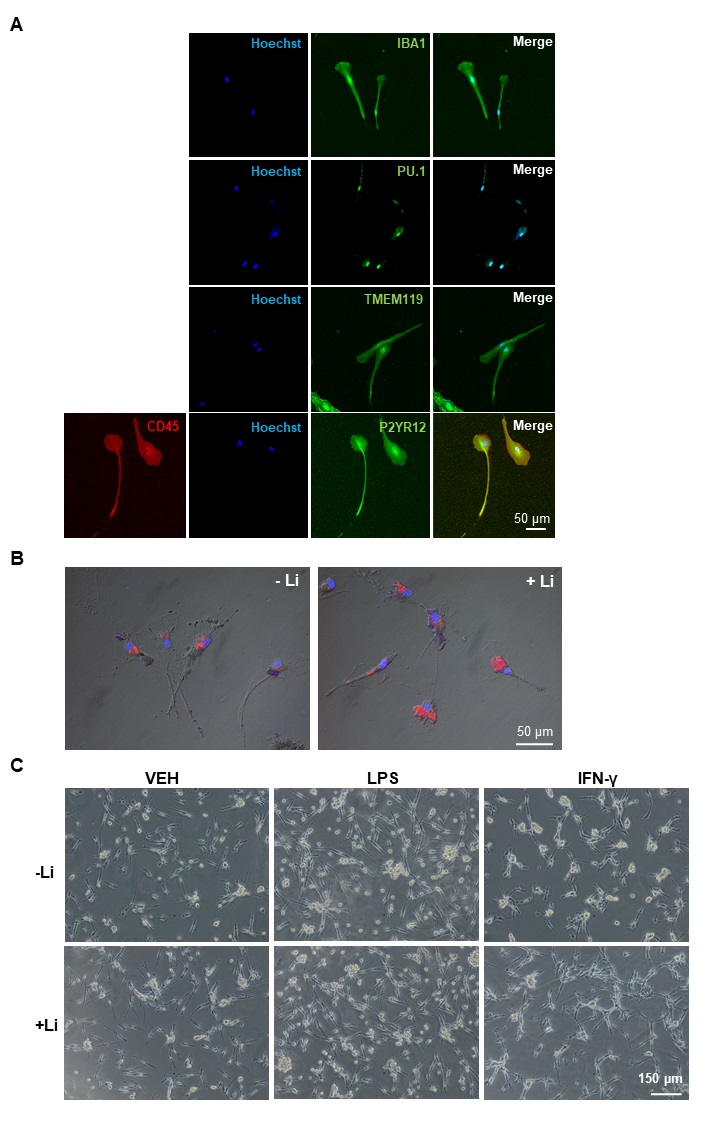
