## Supplementary material for "Lithium inhibits tryptophan catabolism via the inflammation-induced kynurenine pathway in human microglia": Table S3

| REAGENT or RESOURCE | SOURCE | IDENTIFIER |
| --- | --- | --- |
| **Antibodies** | | |
| Alexa488 | Thermo Fisher Scientific | Cat#A21202 RRID: AB_141607, Cat#A21206 RRID: AB_2535792 |
| Alexa647 | Thermo Fisher Scientific | Cat#A31573 RRID: AB_2536183 |
| APC anti-human CD43 | BioLegend | Cat#343205  RRID: AB_2194073 |
| Go-ChIP-grade STAT1 | BioLegend | Cat#686413  RRID: AB_2687272 |
| Go-ChIP-grade STAT3 | BioLegend | Cat#651015  RRID: AB_2715801 |
| Mouse anti-ACTB | Abcam | Cat#8226 |
| Mouse anti-CD45 | EXBIO | Cat#11-222-C025, Clone ID: MEM-28 |
| Mouse anti-GSK-3β (3D10) | Cell Signaling Technology | Cat#9832 |
| Mouse anti-STAT1 (9H2) | Cell Signaling Technology | Cat#9176 |
| Mouse anti-STAT3 (124H6) | Cell Signaling Technology | Cat#9139 |
| Rabbit anti-GAPDH (14C10) | Cell Signaling Technology | Cat#2118 |
| Rabbit anti-IBA1 | Wako Chemicals | Cat#019-19741 |
| Rabbit anti-IDO1 | antibodies-online | Cat#ABIN1169196 |
| Rabbit anti-IDO1 (D5J4E™) | Cell Signaling Technology | Cat#86630 |
| Rabbit anti-P2RY12 | Biozol | Cat#HPA014518 |
| Rabbit anti-phospho-GSK3β (Ser9) (D85E12) | Cell Signaling Technology | Cat#5558 |
| Rabbit anti-phospho-STAT1 (Ser727) | AAT Bioquest | Cat#8A7221 |
| Rabbit anti-phospho-STAT3 (Tyr705) | Cell Signaling Technology | Cat#9131 |
| Rabbit anti-PU.1 | Cell Signaling | Cat#2258 |
| Rabbit anti-TMEM119 | Biozol | Cat#HPA051870 |
| **Biological Samples** | | |
| Human brain cortical biopsy material | Charité-Universitätsmedizin Berlin | N/A |
| **Chemicals, Peptides, and Recombinant Proteins** | | |
| Activin-A | Peprotech | Cat#120-14E |
| B27 | Thermo Fisher Scientific | Cat#17504044 |
| BMP4 | Miltenyi Biotec | Cat#30-111-164 |
| CD200 | Novoprotein | Cat #C311 |
| Chemically-defined lipid concentrate | Thermo Fisher Scientific | Cat#11905-031 |
| CX3CL1 | Peprotech | Cat#300-31 |
| F12 | Thermo Fisher Scientific | Cat#11765054 |
| Fetal calf serum | Biochrom, Merck | Cat#S0613 |
| FGF2 | Peprotech | Cat#100-18B |
| Geltrex | Thermo Fisher Scientific | Cat#A1413302 |
| Glutamax | Thermo Fisher Scientific | Cat#35050061 |
| Hoechst dye 33342 | Abcam | Cat#ab228551 |
| Human IFN-γ | Peprotech | Cat#300-02 |
| IL-3 | Peprotech | Cat#200-03 |
| IL-34 | Peprotech | Cat#200-34 |
| IL-6 | Peprotech | Cat#200-06 |
| IMDM | Thermo Fisher Scientific | Cat#12440053 |
| Insulin | Sigma-Aldrich | Cat#I2643 |
| ITS-G | Thermo Fisher Scientific | Cat#414000045 |
| ITSG-X | Thermo Fisher Scientific | Cat#51500056 |
| L-ascorbic acid 2-phosphate magnesium | Sigma-Aldrich | Cat#A8960 |
| Lithium chloride | Sigma-Aldrich | Cat#L4408 |
| Live cell imaging solution | Thermo Fisher Scientific | Cat#A14291DJ |
| L-kynurenine | Sigma-Aldrich | Cat#K8625 |
| LPS *E.coli* O55: B5 | Sigma-Aldrich | Cat#L2880 |
| L-tryptophan | Sigma-Aldrich | Cat#93659 |
| M-CSF | Peprotech | Cat#300-25 |
| Microglia medium | Innoprot | Cat#P60116 |
| Mobile phase MD-TM | Thermo Fisher Scientific | Cat#70-1332 |
| Monothioglycerol | Sigma-Aldrich | Cat#M1753 |
| N2 | Thermo Fisher Scientific | Cat#17502048 |
| Neural Tissue Dissociation Kit (P) | Miltenyi Biotec | Cat#130-092-628 |
| Nifuroxazide | Sigma-Aldrich | Cat#46494 |
| Non-essential amino acids | Thermo Fisher Scientific | Cat#11140035 |
| PathScan® Cell lysis buffer | Cell Signaling Technology | Cat#7018 |
| Phenol-free DMEM/F12 | Thermo Fisher Scientific | Cat#11039021 |
| PVA | Sigma-Aldrich | Cat#8136 |
| Rock Inhibitor (RI) | Stem Cell Technology | Cat#725252 |
| SB-216763 | AdipoGen | Cat#AG-CR1-3659 |
| SCF | Peprotech | Cat#300-07 |
| TeSR-E8 | Stem Cell Technologies | Cat#05940 |
| TGFb-1 | Peprotech | Cat#100-21 |
| Thiazolyl Blue Tetrazolium Bromide (MTT) | Sigma-Aldrich | Cat#M5655 |
| TPO | Peprotech | Cat#300-18 |
| VEGF | Peprotech | Cat#100-20 |
| **Critical Commercial Assays** | | |
| Imprint® Chromatin Immunoprecipitation kit | Sigma-Aldrich | Cat#CHP1 |
| Mesoscale MSD^©^ Multi-Spot Assay System | Mesoscale | Cat#K15008B |
| PathScan®Immune antibody array kit | Cell Signaling Technology | Cat#13788 |
| pHrodo Red S. aureus | Thermo Fisher Scientific | Cat#A10010 |
| **Experimental Models: Cell Lines** | | |
| BIHi001-A | Berlin Institute of Health (BIH) | N/A |
| BIHi004-A | Berlin Institute of Health (BIH) | N/A |
| BIHi005-A | Berlin Institute of Health (BIH) | N/A |
| Immortalized human microglia | Innoprot | Cat#P10354-IM |
| **Oligonucleotides** | | |
| Primer for *IDO1*  Forward ctgttccttactgccaactctc  Reverse cgtccatgttctcataagtcagg | This paper | N/A |
| Primer for *IDO2*  Forward cagtgccattgtctttggaaagc  Reverse cttgaagctggtgagcatcaac | This paper | N/A |
| Primer for *REEP5*  Forward gaactgcatgactga ccttctg  Reverse cagcaatgctgaacacaccatac | This paper | N/A |
| Primer for *region2_ChIP*  Forward TCATGTAGTCATAAGAAACACAGTC  Reverse GCATATGGCTTTCGTTACAGTCT | This paper | N/A |
| Primer for *TDO*  Forward ctatgggaactacctgcatttg  Reverse ccagaggatttgcttaaaccag | This paper | N/A |
| **Software and Algorithms** | | |
| Chromeleon™ 7.2 Chromatography Data System (CDS) software | Thermo Fisher Scientific | RRID:SCR_016874 |
| GraphPad Prism 8 | GraphPad | RRID:SCR_002798 |
| LI-COR® Image Studio array analysis software | LI-COR | RRID:SCR_015795 |
| Zen® software | Carl Zeiss | RRID:SCR_018163 |
| **Other** | | |
| Spin-X® centrifuge tube filter | Corning | Cat#8161 |
